# Phosphorylation alters FMRP granules and determines their transport or protein synthesis abilities

**DOI:** 10.1101/2023.03.15.532613

**Authors:** Shivani C. Kharod, Dong-woo Hwang, Heejun Choi, Kyle J. Yoon, Pablo E. Castillo, Robert H. Singer, Young J. Yoon

**Author notes:** Correspondence Young J. Yoon, Ph.D. Dominick P. Purpura Department of Neuroscience Albert Einstein College of Medicine 1300 Morris Park Avenue Bronx, NY 10461 USA.

## Abstract

Fragile X messenger ribonucleoprotein (FMRP) is an RNA-binding protein implicated in autism that suppresses translation and forms granules. While FMRP function has been well-studied, how phosphorylation regulates granule binding and function remains limited. Here, we found that Fragile X patient-derived I304N mutant FMRP could not stably bind granules, underscoring the essential nature of FMRP granule association for function. Next, phosphorylation on serine 499 (S499) led to differences in puncta size, intensity, contrast, and transport as shown by phospho-deficient (S499A) and phospho-mimic (S499D) mutant FMRP granules. Additionally, S499D exchanged slowly on granules relative to S499A, suggesting that phosphorylated FMRP can attenuate translation. Furthermore, the S499A mutant enhanced translation in presynaptic boutons of the mouse hippocampus. Thus, the phospho-state of FMRP altered the structure of individual granules with changes in transport and translation to achieve spatiotemporal regulation of local protein synthesis.

**Teaser:** The phosphorylation-state of S499 on FMRP can change FMRP granule structure and function to facilitate processive transport or local protein synthesis.

## MAIN TEXT

## Introduction

Fragile X messenger ribonucleoprotein (FMRP) is an RNA-binding protein implicated in mRNA transport and protein synthesis (*1–3*). Loss of expression or loss-of-function mutations can lead to intellectual disability with links to autism (*4*). In *Fmr1* knock-out mice, the absence of FMRP led to elevated protein production and altered synaptic plasticity (*5, 6*). At the molecular level, studies have shown that FMRP interacts with translating polyribosomes (*7, 8*) and mRNA (*9, 10*) to regulate protein synthesis and travel along dendrites in an activity-dependent manner (*11, 12*). In addition, recent studies have elucidated different mechanisms of translational repression by FMRP through interactions with the FMRP-interacting factor, CYFIP1 (*13*), stalled ribosomes (*10*) and microRNA complexes (*14*). In mice, FMRP is phosphorylated on serine 499 (S499) which enhances its association with polysomes and stalled ribosomes (*15*). Subsequent studies reported that activation of metabotropic glutamate receptors (mGluR) led to brief dephosphorylation of FMRP by protein phosphatase 2A (PP2A) (*16*) and facilitated the local translation of *Arc* mRNA in dendrites (*17*). Also, dephosphorylated FMRP gets ubiquitinated for degradation by the proteasome (*18*). More recently, we reported that FMRP phosphorylation plays a critical role in presynaptic structural and functional plasticity (*19*). Altogether, these studies point to phosphorylation of S499 as a critical determinant in regulating FMRP function in translation.

One prominent feature of FMRP in neurons is the formation of Fragile X granules (*12, 20–22*). FMRP is a multivalent protein that can interact with mRNA, ribosomes, and other regulatory proteins for transport and translational control in dendrites (*2, 23*). FMRP can also bind with itself to mediate self-assembly into relatively large intracellular structures in vitro (*24, 25*). However, how changes in FMRP granule structure are linked to changes in FMRP function in living cells remains unresolved. For instance, it is unclear whether the Fragile X patient-derived I304N FMRP mutation, which has lost the ability to bind ribosomes, can still form granules. More importantly, how phosphorylation and dephosphorylation of FMRP can modify the structure of FMRP granules is unexplored. By uncovering the features of FMRP granules that are translation-permissive or translation-repressive, we can predict and assign the function of granules based on their structure and motion.

Here, we have investigated the granule-binding ability of GFP-FMRP, S499A, S499D, and I304N to understand their impact on granule structure and function. Our findings suggest that constitutively phosphorylated S499D mutant possesses greater processivity and long-distance travel of FMRP granules by slowing translation. In contrast, S499A granules may favor scanning through the dendrite through intermittent pauses to deliver newly synthesized proteins at synapses. Thus, S499 phosphorylation is critical to changes in FMRP granule structure and may function as a phospho-switch to regulate transport and local protein synthesis.

## Results

### Patient-derived I304N mutant FMRP does not stably bind FMRP granules

A rare missense mutation on FMRP, I304N, was originally discovered in a Fragile X patient who exhibited severe autistic behavior but expressed normal levels of FMRP mRNA and protein (*26*). Sequencing the *Fmr1* locus revealed that the patient had a missense mutation within the KH2 domain of FMRP. Moreover, I304N mutant FMRP failed to co-fractionate with polyribosomes in an I304N patient-derived cell line (*27*) and the I304N mouse model (*28*). We wanted to see if the loss of polysome interaction could affect its ability to bind granules. When we expressed the I304N FMRP (GFP-I304N) in neurons, we observed very few discrete puncta along with diffuse fluorescence within dendrites (Fig. 1A). Continuous imaging of GFP-I304N in dendrites showed no discernible granules over time in kymographs (Fig. 1B), indicating that either it was unable to bind FMRP granules or that the bound population was obscured by the majority unbound population. To achieve a higher resolution view of I304N FMRP movement, we conducted single molecule tracking of individual FMRP proteins using HaloTag-I304N FMRP (Fig. 1, C and D). HaloTag uses self-labeling technology where bright, cell-permeable dyes conjugated to a HaloTag ligand can covalently bind to the HaloTag, thereby rendering single molecules of Halo-FMRP visible by fluorescence microscopy (*29*). We used deuterated JFX dyes (*30*), which have improved photostability and ideal for long-term single molecule imaging. When we compared the movement of individual Halo-FMRP and Halo-I304N molecules, the trajectories for Halo-FMRP were either stationary or followed a linear path (Fig. 1C and movie S1), suggesting that the moving FMRP proteins were traveling along the cytoskeleton consistent with active transport. In contrast, most Halo-I304N molecules appeared untethered with a large degree of freedom in their movement (Fig. 1D and movie S1). On average, I304N FMRP proteins had higher median displacement and velocity than Halo-FMRP (Fig. 1, E and F). By plotting the particle intensity and size we confirmed that no changes or loss of signal were observed during image acquisition due to increased speed or photosensitivity (Fig. 1, G and H). These results suggested that while I304N FMRP had not completely lost affinity for molecular interactions, the binding time was so brief that the unbound state prevailed where most proteins appeared to be freely diffusing. Notably, the lack of stable granule-binding by the I304N FMRP would indicate that its interaction with ribosomes is essential for FMRP granules, and this loss of FMRP on granules is linked to FXS pathophysiology.

**Fig. 1.**
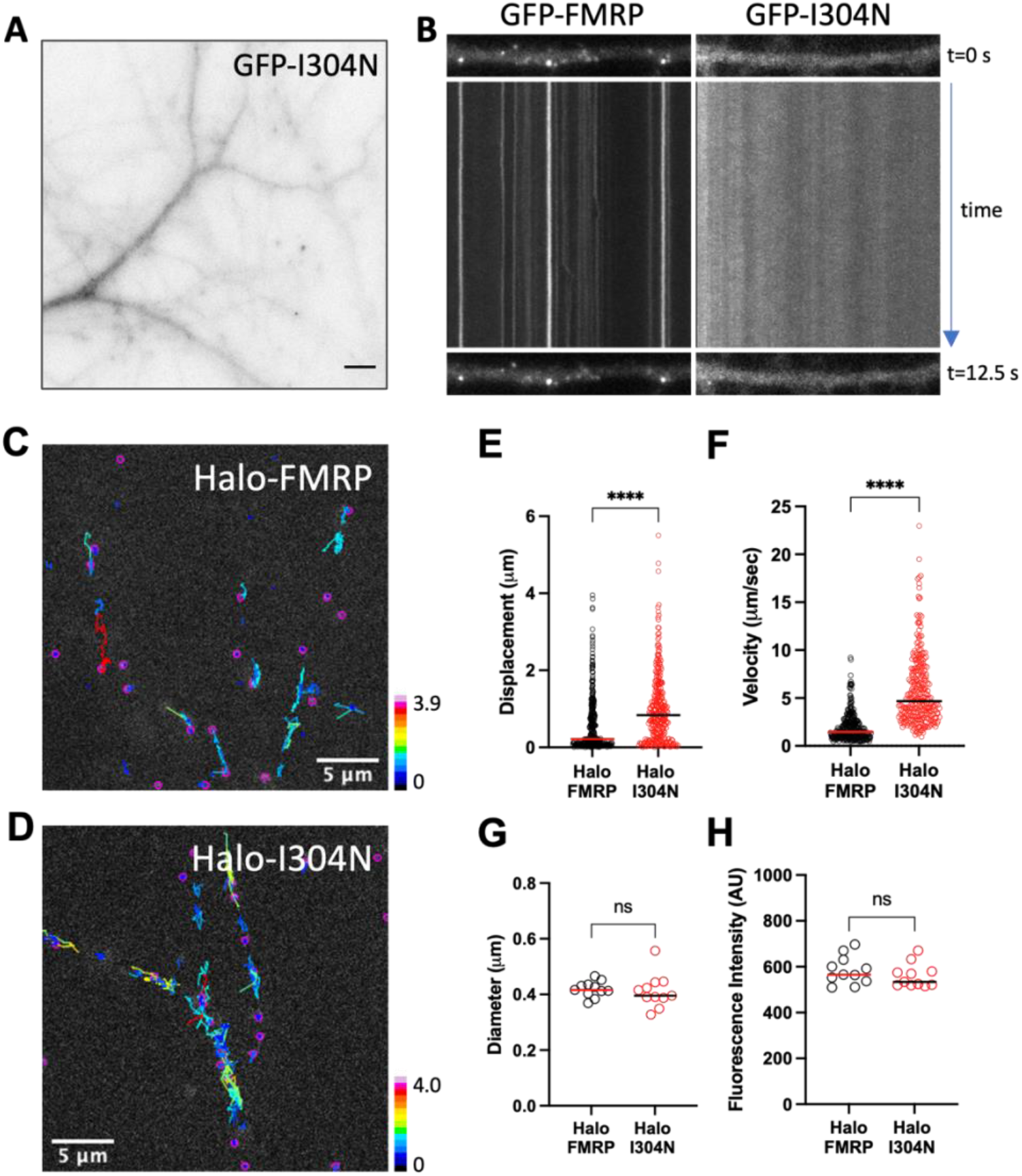
FXS patient-derived I304N FMRP does not stably bind FMRP granules. (**A**) Representative fluorescence image (pseudo-colored as inverted grayscale) of a dendrite expressing the I304N mutant FMRP for one week. Scale bar is 5 μm. (**B**) Comparison of kymographs from GFP-FMRP (left) or GFP-I304N (right) expressing dendrites. The discrete stationary fluorescent puncta present in GFP-FMRP are depicted as continuous lines along the time-axis while the I304N dendrite does not exhibit any stationary particles. (**C**) Sparse labeled HaloTag-FMRP (Halo-FMRP, pink circles) overlaid with particle trajectories that are color-coded by total displacement. LUT denotes total displacement during 5 second imaging epoch ranging from 0 to 3.9 μm. Scale bar is 5 μm. See movie S1. (**D**) Sparse labeled HaloTag-I304N FMRP (Halo-I304N, pink circles) overlaid with particle trajectories that are color-coded by total displacement. See movie S1. LUT denotes total displacement during 5 second imaging epoch ranging from 0 to 4.0 μm. (**E-F**) Scatter plot of total displacement or mean velocity of Halo-FMRP (black circles; n=359 tracks from 11 cells) or Halo-I304N (red circles; n=299 tracks from 11 cells) particles. The horizontal lines denote population median. Statistical significance was tested using unpaired, two-tailed Mann-Whitney test. p**** < 0.0001. (**G-H**) Scatter plot of Halo-FMRP (n=11) and Halo-I304N particles (n=11) mean particle diameter and intensity per cell. The horizontal lines denote population mean. Statistical significance was tested using unpaired, two-tailed t-test. ns, not significant.

### S499 phospho-mutant FMRP form distinct granules

From activity-dependent dephosphorylation (*16, 17*) to hierarchical phosphorylation of neighboring serine residues (*15, 31*), S499 has been implicated as the major phospho-switch that governs FMRP function (*23, 32*). We reasoned that mutations on S499 that block or mimic phosphorylation would allow us to capture intermediates of FMRP granules. We generated N-terminal GFP fusions to wildtype and phospho-mutant FMRP (*15*) driven by the human Synapsin promoter (Fig. 2A). FMRP reporters were expressed in cultured hippocampal neurons at DIV7 for one week to assemble with endogenous FMRP granules. As FMRP granules contain more than one FMRP (*21*), reporter expression results in exchange with FMRP in granules and becomes fluorescent over time (Fig. 2, B and C). To confirm incorporation into FMRP granules, we performed immunofluorescence to GFP-FMRP and endogenous FMRP in dendrites and observed colocalization (fig. S1A). Moreover, co-expression of GFP-FMRP with Halo-FMRP (*19*) or ribosomal protein L10A fused to HaloTag showed colocalization between FMRP with different tags and with ribosomes, respectively (fig. S1, B and C), demonstrating that our FMRP reporters can assemble onto FMRP granules. As neurons are sensitive to levels of FMRP (*33*), we expressed our reporters for one week in immature neurons (DIV7-14). Dendritic expression levels of GFP-FMRP and the phospho-mimic (S499D) mutant were similar, whereas the phospho-deficient (S499A) mutant was reduced as a result of degradation by the proteasome (*18*) (fig. S1D). The levels of expression were ideal to measure differences in FMRP granules (Fig. 2, D and E), as shorter expression (DIV7-10) did not reveal significant differences among the reporters (fig. S1, E to G).

**Fig. 2.**
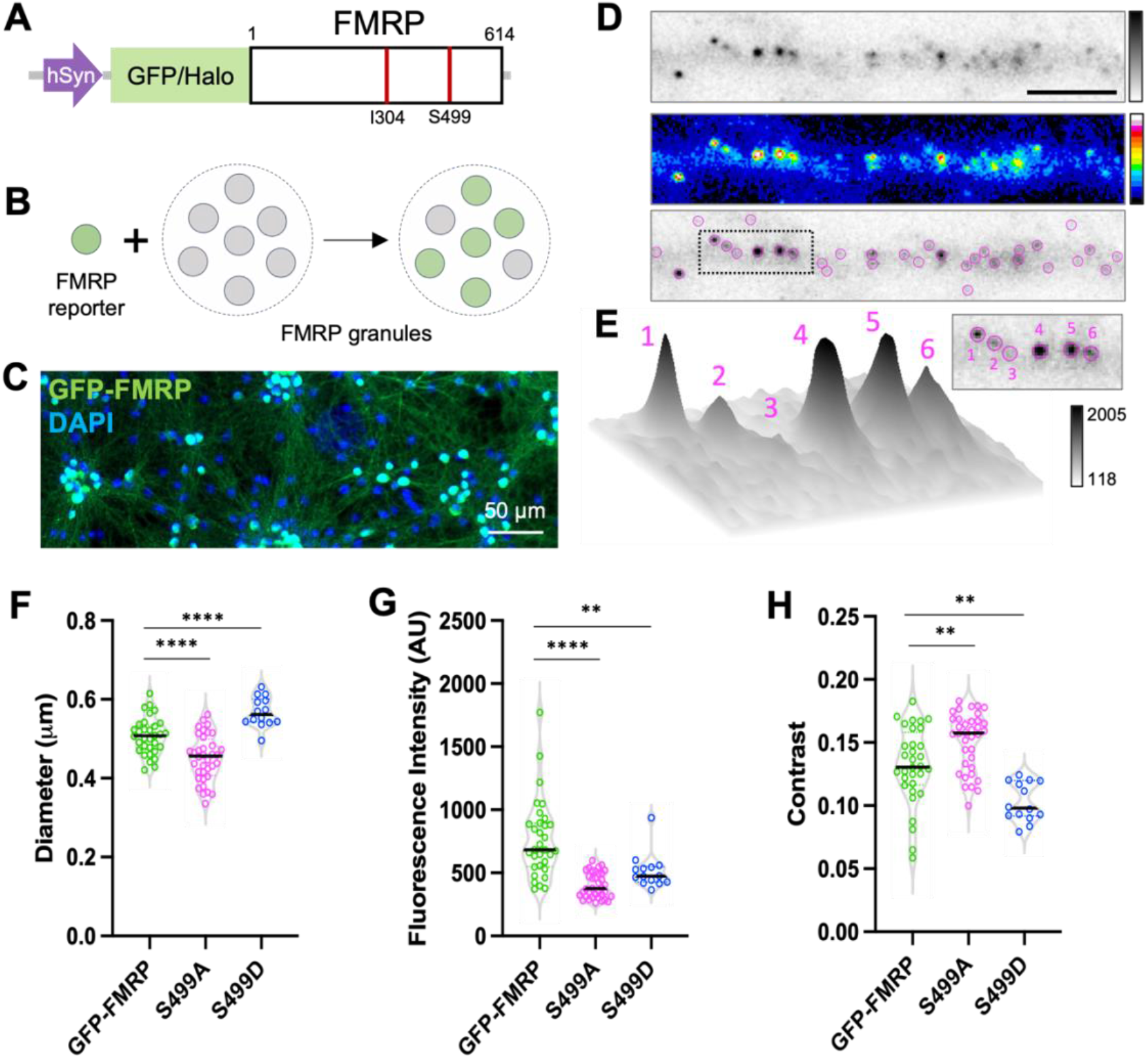
S499 phospho-mutant FMRP form distinct granules. (**A**) Schematic of fluorescent FMRP reporter constructs: human Synapsin promoter drives the expression of GFP-FMRP. Red lines denote location of I304 and S499. (**B**) Cartoon of FMRP reporter in green binding to FMRP granules. Over time, fluorescent FMRP exchanges with endogenous FMRP which allows granules to become fluorescent. (**C**) 10x magnification view of cultured neurons expressing GFP-FMRP labeled with antibodies to GFP shown in green. DAPI is shown in blue. Scale bar is 50 μm. (**D**) Representative dendrite expressing GFP-FMRP (top panel, inverted grayscale). Same dendrite shown in 16-color LUT (middle panel). Overlay with TrackMate particle detection (pink circles, bottom panel). Scale bar is 5 μm. (**E**) Surface plot of FMRP puncta shown in the dotted box in **D**. Numbers on surface plot correspond to puncta in inset. The LUT indicates the minimum and maximum fluorescence intensity values. (**F-H**) Violin plots of granule diameter, fluorescence intensity, or contrast from GFP-FMRP (green; n=32; 10905 granules), S499A (magenta; n=36; 4923 granules) and S499D (blue; n=14; 3088 granules). Horizontal lines denote population median. Statistical significance was calculated using unpaired, two-tailed student’s t-test. p** < 0.01; p**** < 0.0001.

First, we characterized FMRP granules bound by our FMRP reporters to elucidate size, intensity, and contrast as indicators of granule dimensions, amount of FMRP molecules present in granules, and how well each reporter can cluster (*34*), respectively (Fig. 2, F to H). The median diameter of GFP-FMRP granules was around 0.5 μm, consistent with previous reports (*21*). The fluorescence intensity of GFP-FMRP granules exhibited a large distribution, suggesting that cycles of FMRP phosphorylation and dephosphorylation can facilitate more FMRP molecules to incorporate into individual granules. Notably, GFP-FMRP granules showed a range of contrast values encompassing the phospho-mutants, suggesting a mixed population of granules with either predominantly phosphorylated or dephosphorylated FMRP granules. The S499A mutant formed granules that were relatively lower in size and intensity, suggesting that these granules were small but tightly clustered (fig. S2A). The higher contrast value can be attributed to S499A being ubiquitinated and degraded leading to reduced overall dendritic levels (*18*). Conversely, the S499D mutant formed larger granules with slightly higher intensity and lower contrast than S499A, indicating that S499D is bound diffusely on FMRP granules, giving a less compact appearance (fig. S2B). Since S499D mutant had been suggested to bind stalled polysomes preferentially (*15*), S499D-containing granules could be more disordered than a translating granule.

### FMRP phospho-mutant reporters differ in processive movement

As FMRP granules can travel throughout neurons (*11, 35, 36*), we next asked whether FMRP mutants vary in their transport in dendrites. We acquired short continuous images (50 ms per frame for 400 frames) of dendrites expressing FMRP reporters (Fig. 3A) and used TrackMate (LAP tracker) to analyze and classify moving particles (fig. S3, A to C and movie S2 to S4) based on processivity. Processivity of transport, or continuous activity of molecular motors before detaching, is a measure of how efficiently intracellular cargo can travel to discrete locations inside cells. In our analysis, we defined processive motion as a particle that moves continuously along the same direction for five or more consecutive time points, where each processive particle was sorted by its total displacement. We observed two populations of processive GFP-FMRP granules with high and low displacements (Fig. 3B). The low displacement population was prominent in dephosphorylated S499A granules, suggesting they tended to travel shorter distances with intermittent pauses or changes in direction, consistent with scanning along the dendrite. On the other hand, phosphorylated S499D granules showed another population with higher displacements indicative of longer travel distances during each processive motion. These results suggest that phosphorylated FMRP granules favor long-distance transport while dephosphorylated FMRP granules prefer stop-and-go processive motion in search of a docking site.

**Fig. 3.**
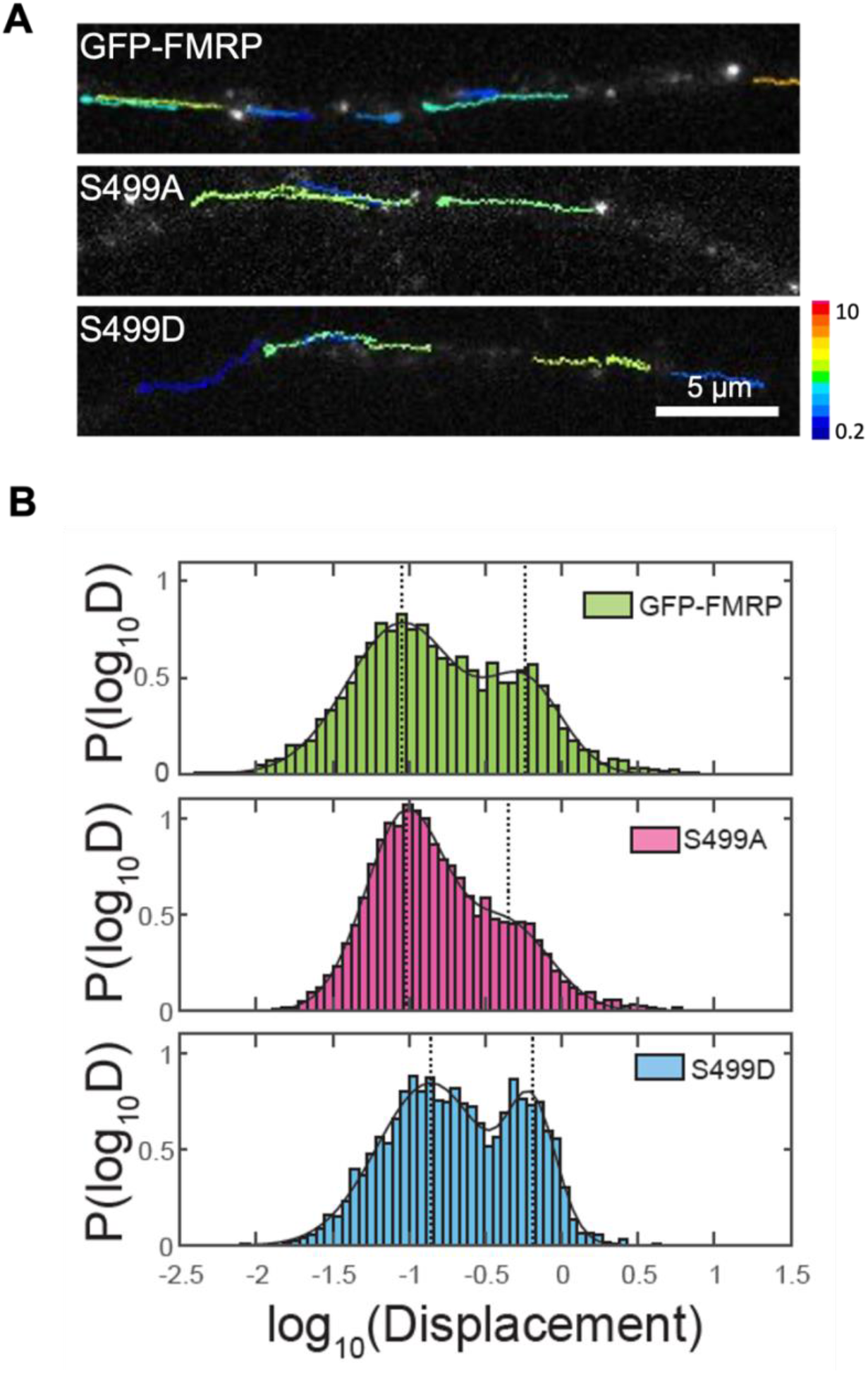
FMRP phospho-mutant granules differ in processive movement. (**A**) Fluorescence images of GFP-FMRP (top), S499A (middle) and S499D (bottom) expressing dendrites overlaid with tracks detected by TrackMate. LUT indicates velocity at μm/sec and the scale bar is 5 μm. See movies S2-S4. (**B**) Histogram of log_10_(Displacement) of GFP-FMRP (top, green), S499A (middle, magenta), and S499D (bottom, blue) granules where displacement is calculated as the distance from the beginning and end of processive movement in microns. Histograms were normalized by the total counts in the population. Dotted lines indicate two peaks from distributions. For GFP-FMRP, P_<HIGH displacement>_ = 0.366 and P_<LOW displacement>_ = 0.634; Displacement_high_ = 0.58 μm and Displacement_low_ = 0.09 μm. For S499A, P_<HIGH displacement>_ = 0.30 and P_<LOW displacement>_ = 0.70; Displacement_high_ = 0.45 μm and Displacement_low_ = 0.09 μm. For S499D, P_<HIGH displacement>_ = 0.436 and P_<LOW displacement>_ = 0.564; Displacement_high_ = 0.64 μm and Displacement_low_ = 0.14 μm.

### FMRP exchange onto FMRP granules requires translation

Given that FMRP phospho-mutants differ in their ability to associate with granules, this suggested that FMRP granules were not static structures. We conducted fluorescence recovery after photobleaching (FRAP) on individual FMRP granules to test whether unbound FMRP proteins could exchange onto FMRP granules. We used low-power focused UV light to selectively photobleach a single fluorescent FMRP puncta (Fig. 4A), reaching an average photobleaching efficiency of 85%. As we did not detect rapid recovery within the first minute, we acquired longer time-lapse images at 30-second intervals for 20 minutes. The FRAP assay was performed at room temperature (22 °C) to minimize movement of the photobleached granule and to reduce the likelihood of interference from other granules that actively traffic through dendrites at physiological temperature.

**Fig. 4.**
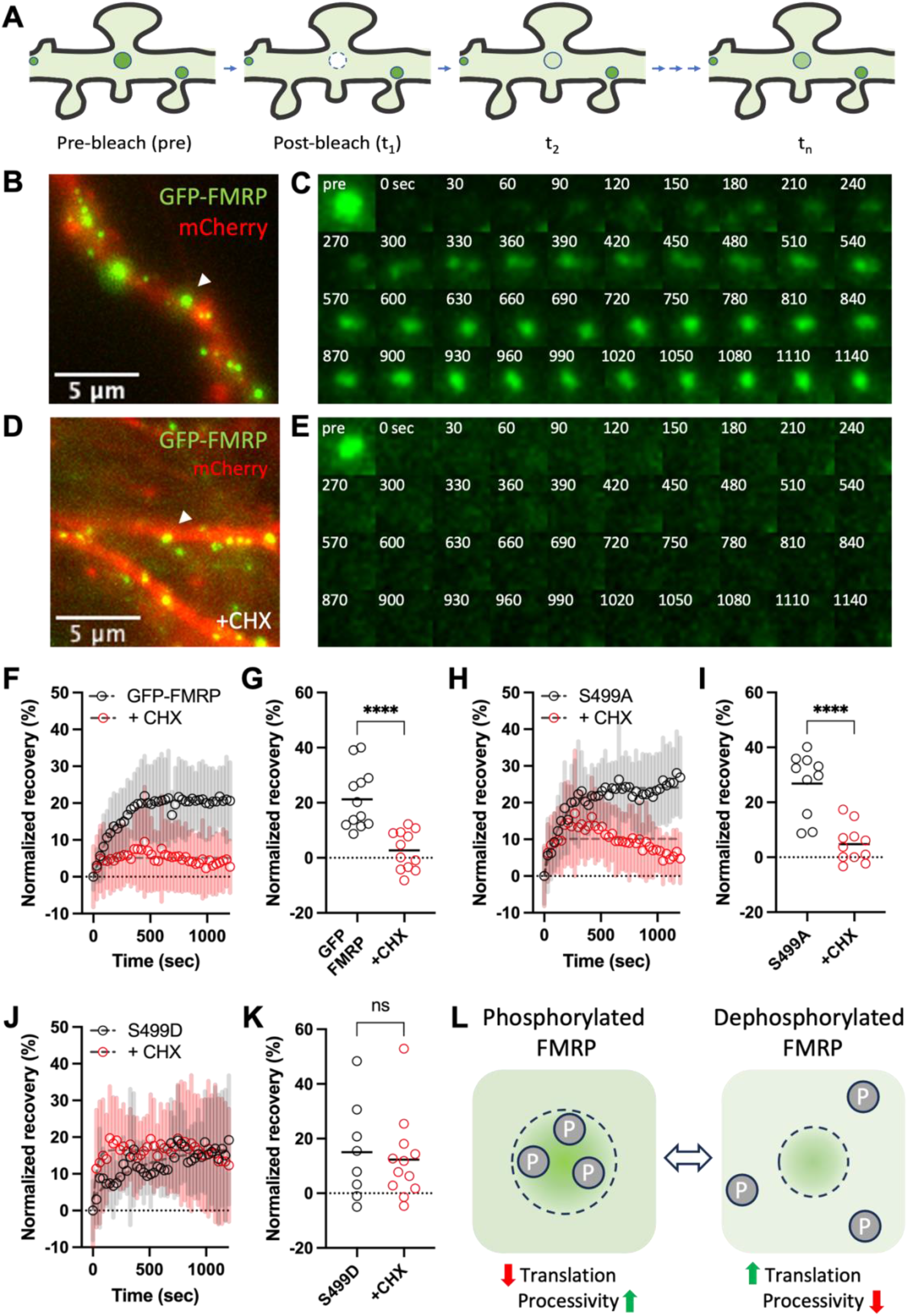
FMRP granules exchange FMRP proteins in a translation-dependent manner. (**A**) Schematic of spot photobleaching of individual FMRP granules. (**B**) Representative image of GFP-FMRP granule (green) in dendritic segment (red) selected for the fluorescence recovery assay. The white arrowhead indicates the photobleached granule and the scale bar is 5 μm. See movie S5. (**C**) Time-series montage of FMRP granule recovery. Time is noted on the top-right in seconds. (**D**) Representative image of GFP-FMRP granule (green) FRAP in the presence of cycloheximide (CHX). The white arrowhead indicates the photobleached granule and the scale bar is 5 μm. See movie S7. (**E**) Time-series montage of FMRP granule recovery in CHX. (**F**) Plot of normalized mean fluorescence recovery ± SD (shaded bars) for granules expressing GFP-FMRP (black circles; n=12) or CHX-treatment (red circles; n=12) over time. The recovery values were fit to a nonlinear regression curve (gray dashed lines) to calculate time constants (τ). The dotted line indicates the average baseline intensity following photobleaching set to zero. (**G**) Normalized recovery of individual granules at the final timepoint for GFP-FMRP (black circles; n=12) or CHX-treatment (red circles; n=12). Horizontal lines denote population mean. Statistical significance was calculated using unpaired, two-tailed student’s t-test. p**** < 0.0001. (**H**) Normalized mean recovery ± SD for S499A granules (black circles; n=10) or CHX-treatment (red circles; n=11). (**I**) Normalized recovery of individual granules at the final timepoint for S499A (black circles; n=10) or CHX-treatment (red circles; n=11). Statistical significance was calculated using unpaired, two-tailed student’s t-test. p**** < 0.0001. (**J**) Normalized mean recovery ± SD for S499D granules (black circles; n=8) or CHX-treatment (red circles; n=12). (**K**) Normalized recovery of individual granules at the final timepoint for S499D (black circles; n=8) or CHX-treatment (red circles; n=12). Statistical significance was calculated using unpaired, two-tailed student’s t-test. ns, not significant. (**L**) Model of how phosphorylation changes FMRP granule dimensions, translation and transport. Dotted circle indicates FMRP granule formed by predominantly phosphorylated (left) or dephosphorylated (right) FMRP. The bidirectional arrow indicates the change in the phospho-state of FMRP. Circled P indicates phosphate groups. Green up-arrow indicates increase and red down-arrow indicates decrease.

Intriguingly, we observed a gradual and partial fluorescence recovery of GFP-FMRP (Fig. 4, B and C and movie S5). When the results were fit to a nonlinear regression curve, GFP-FMRP reached half-maximal recovery after 2.5 minutes (τ = 172.5 s) with an average recovery of 20% after 10 minutes (Fig. 4F; black circles). Also, the recovery plateaus after 10 minutes, which suggested a steady-state exchange between inside and outside of the granule. In contrast, G3BP1-GFP, a well-characterized granule protein (*37, 38*), reached 60% recovery after one minute (fig. S4, A to C and movie S6). To test whether FMRP exchange depended on translation, we performed FRAP in the presence of a translation elongation inhibitor, cycloheximide (CHX), which would result in completely stalled ribosomes. In the presence of CHX, GFP-FMRP granules exhibited a markedly reduced recovery (Fig. 4, D and E and movie S7), suggesting that FMRP bound to CHX-stalled ribosomes do not exchange (Fig. 4F; red circles). Comparison of recovery at the final timepoint showed a significant difference in fluorescence (Fig. 4G). Use of another translation elongation inhibitor, anisomycin, resulted in a similar reduction in recovery (fig. S4 D to G and movie S8). Like GFP-FMRP, recovery of S499A reached 20% (Fig. 4, H and I, fig. S5 A and B and movie S9 and S10) with slightly faster kinetics (τ = 161.0 s). However, in the presence of CHX, the recovered signal gradually diminished, suggesting that S499A has a reduced affinity for stalled ribosomes in granules. S499D fluorescence was the slowest to recover (τ = 256.5 s), eventually reaching 20% recovery after 20 minutes (Fig. 4,J and K, fig. S5, C and D and movie S11 and S12). The recovery is consistent with S499D binding to attenuate actively translating ribosomes and reduce the rate of exchange. Surprisingly, the initial recovery kinetics of S499D was enhanced by CHX-treatment (τ = 30.8 s). As CHX binds to the E-site of the translating ribosome, the stable interaction between phosphorylated FMRP and ribosomes may be occluded, leading to a faster exchange. Taken together, our data show that GFP-FMRP on granules exchange at a timescale consistent with translation, and this exchange is sensitive to translation inhibitors. Moreover, dephosphorylated FMRP preferentially bind translating ribosomes and travel short distances, whereas phosphorylated FMRP exchanges slowly due to attenuating translation while displacing larger distances (Fig. 4L).

Identifying the C-terminal portion of FMRP as a low-complexity region (LCR) was intriguing as it hinted that FMRP granules may have functional consequences (*39*). As FMRP has multiple RNA-binding domains, such as KH domains and RGG-box, it was suggestive that FMRP may undergo phase separation (*40*). We generated a C-terminal truncation mutant, FMRP^1-444^ (ΔC-term), which contained the Agenet and KH1/2 domains and the nuclear export signal but not the S499 and the RGG box to test whether it could bind to granules. When expressed in neurons, ΔC-term FMRP was able to bind granules, albeit less clustered (fig. S6A) as indicated by the large diameter, reduced intensity and contrast similar to G3BP1 (fig S6, B to E), which has been shown to phase separate (*41*). These results support the conclusion that S499 is necessary but not sufficient for tight granule binding. Fluorescence recovery of ΔC-term FMRP granules was also similar to G3BP1 with relatively fast recovery kinetics (τ = 15.2 s), and CHX-treatment had no effect on exchange (fig. S6, F to H and movie S13 and S14). Conversely, when we expressed the C-terminal portion of FMRP^424-614^ (FMRP C-term), we did not detect granules or localization to any discrete subcellular structure (fig. S6I). While our results suggest that the C-terminal region of FMRP may not independently form granules, there could be FMRP-interactors on disparate regions of FMRP that may trigger phase separation (*42, 43*).

### Phospho-deficient FMRP enhances local translation in mossy fiber boutons

Recently, we have shown that presynaptic boutons can perform increased local protein synthesis in response to presynaptic activity in an FMRP-dependent manner (*19, 44*). Presynaptic compartments such as hippocampal mossy fiber boutons provide an ideal subcellular compartment to observe discrete, measurable differences in protein synthesis. To determine whether the phospho-state of FMRP can change the translational output in presynaptic boutons, we injected lentiviruses encoding Halo-FMRP or Halo-S499A into the dentate gyrus region of the mouse hippocampus to target dentate granule cells, which give rise to mossy fibers. Following the expression of these constructs, acute hippocampal slices were fluorescently labeled, and mossy fiber boutons were imaged (Fig. 5, A to C). Consistent with our findings in cultured neurons, we found that the S499A mutant formed granules that were smaller, less bright, and more compact when compared to Halo-FMRP (Fig. 5, D to F). Next, to determine whether the difference in granule clustering led to causal changes in protein synthesis, we used puromycin (PMY) incorporation onto nascent peptides (*45*) to measure the total output of protein synthesis in the mossy fiber tract (Fig. 5, G and H). Using FMRP fluorescence as a mask, we quantified the average intensity of the colocalized puromycin label and found that S499A granules correlated with an increase in protein synthesis when compared to Halo-FMRP (Fig. 5I). Our results demonstrate that regulation of FMRP granule structure and protein synthesis in mossy fiber boutons correlates with the phosphorylation state of FMRP.

**Fig. 5.**
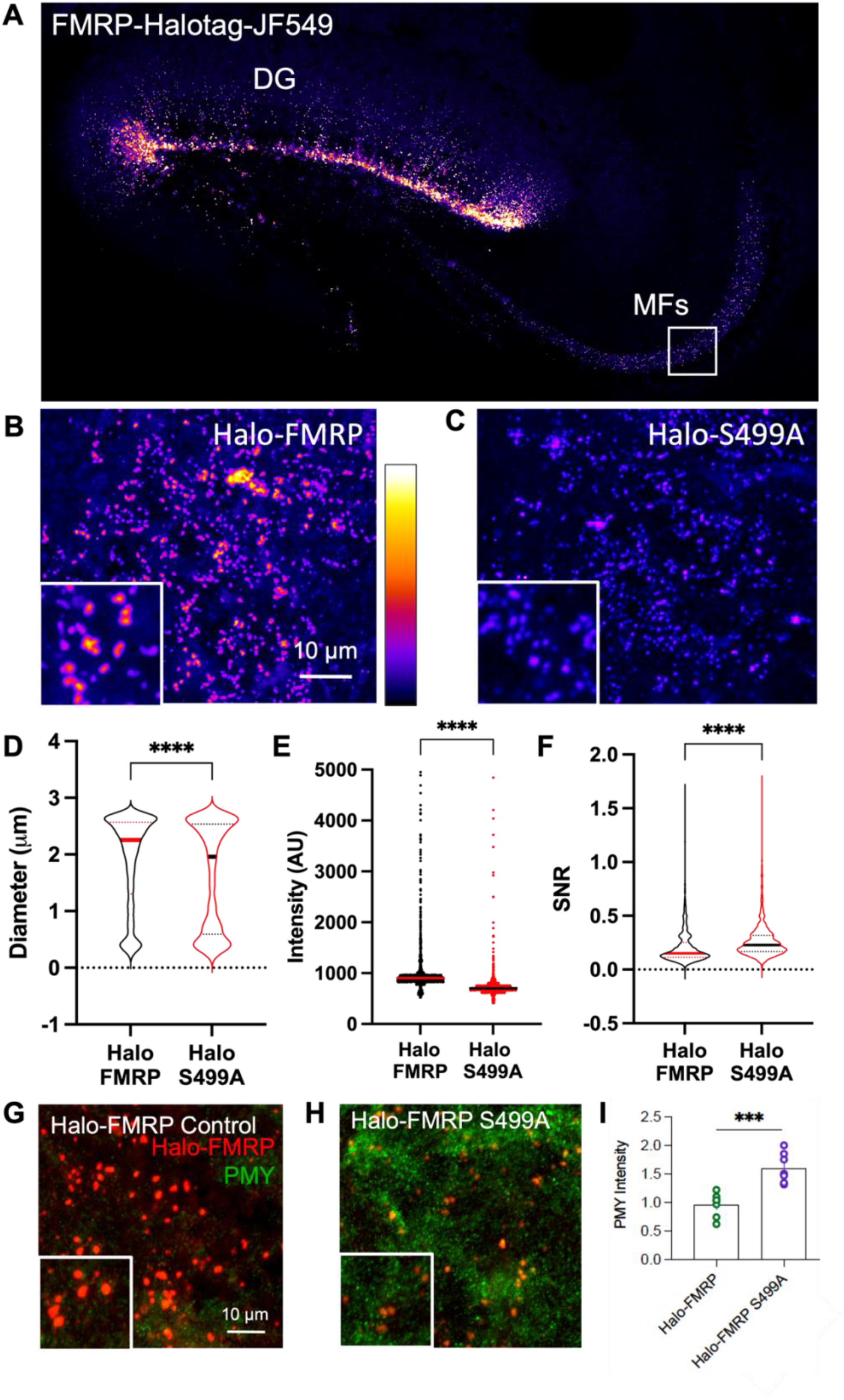
Halo-S499A FMRP leads to increased protein synthesis in mossy fiber boutons. (**A**) Representative tiled fluorescence image of a hippocampal slice expressing Halo-FMRP labeled with JF549 HaloTag ligand after targeted stereotactic injection into the dentate gyrus (DG). Mossy fiber projections (MFs) containing boutons from Halo-FMRP-expressing neurons are shown in the white box. (**B-C**) Representative images of mossy fiber boutons expressing Halo-FMRP or Halo-S499A. Scale bar is 10 μm. Insets show higher magnification of boutons. (**D-F**) Violin plot of Halo-FMRP (black; n=3650 boutons) and Halo-S499A (red; n=2556 boutons) granule diameters, intensities, and signal-to-noise ratio (SNR). Horizontal lines denote population median. Statistical significance was calculated using unpaired, two-tailed student’s t-test. p**** < 0.0001. (**G-H**) Representative images of puromycin (PMY, green) and JF549 HaloTag labeling (red) in slices from mice injected with Halo-FMRP (left) or Halo-S499A (right). FMRP fluorescence was used as a mask to measure puromycin signal. Scale bar is 10 μm. Insets show higher magnification of boutons. (**I**) Average intensity of colocalized puromycin label in Halo-FMRP (n=6 slices) and Halo-S499A (n=6 slices) were normalized to Halo-FMRP. Statistical significance was calculated using unpaired, two-tailed student’s t-test. p**** < 0.001.

## Discussion

In this study, we set out to better understand FMRP granule structure and function. Using FMRP reporters and high resolution imaging in neurons harboring endogenous FMRP granules helped identify distinguishing features that differ in mutant FMRP. For instance, our observation that the FXS patient-derived I304N mutation could not form discrete granules suggests a possible linkage to FXS. The integrity and maintenance of FMRP on granules could be an important factor in neuronal function and physiology.

To capture FMRP granule intermediates, we focused on the well-characterized phospho-switch S499 of FMRP. The current model of FMRP function is that phosphorylation on S499 results in a translation-repressive state, and dephosphorylation leads to a translation-permissive state, where FMRP likely mimics the phosphorylated state in neurons (*15*). Upon mGluR activation, FMRP is transiently dephosphorylated, thereby providing a temporal window of protein synthesis in an activity-dependent manner. Using our FMRP phospho-mutant reporters, we could distinguish differences in FMRP granules that were translation-permissive or -repressive (Fig. 2 and 3). Regarding granule size and contrast, our observations were consistent with our expectations, where the S499A mutant bound smaller and more discrete granules, while the S499D mutant bound larger and less discrete granules relative to GFP-FMRP, reflecting their differing affinities for granules. However, when we looked at the fluorescence intensity of GFP-FMRP granules, on average, they were much brighter than either mutant. By blocking S499 from phosphorylation and subsequent phosphorylation of neighboring serines (*15, 31*), the phospho-mutant FMRP could not be incorporated into granules in numbers comparable to GFP-FMRP. This observation suggests that phosphorylation and dephosphorylation cycles on S499 can promote phosphorylation of neighboring serines such that more GFP-FMRP can bind or assemble onto granules.

Given the differences in granules, we reasoned that the interaction of our FMRP reporters with granules was not a static process. Our FRAP results for GFP-FMRP showed a partial recovery that was blocked by the translation elongation inhibitor, CHX. The observation that GFP-FMRP could exchange on granules through translating ribosomes was inconsistent with the model that FMRP functions predominantly to stall ribosomes. However, it is conceivable that FMRP may be tuning translation to enhance the fidelity of local protein synthesis. For instance, a previous report showed that FMRP binds to its target mRNA along the coding sequence (*10*), suggesting that FMRP-binding was not sequence-specific within the open reading frames of target transcripts. The results do not exclude the possibility that FMRP interaction with the coding regions of mRNA could occur on attenuated or stalled ribosomes. Second, our β-actin translation reporter (which is not an FMRP target) displayed a half-maximal recovery time of around 75 seconds in dendrites (*46*), compared to >150 seconds with our GFP-FMRP reporters (Fig. 4), suggesting that FMRP-binding may be slowing translation. Third, the observation that FMRP occupancy on mRNA can stabilize target transcripts through codon optimization (*47*) will likely require active translation to protect FMRP targets from mRNA surveillance and decay mechanisms (*48*). Taken together, there is a possibility that FMRP may enhance the fidelity of local protein synthesis through attenuation of translation rates rather than completely stalling translation.

Unexpectedly, S499D mutant exhibited an ability to exchange with CHX-stalled FMRP granules (Fig. 4J). This result was surprising as phosphorylated FMRP should preferentially remain bound to stalled ribosomes. The initial rapid recovery or exchange indicated less stable binding to granules due to CHX-induced stalling. One explanation could be that FMRP may prefer to stall translation at a specific elongation step of protein synthesis that “locks-in” FMRP onto granules, and stalling ribosomes with CHX can lead to changes in ribosome conformation, thereby destabilizing this interaction. The cryo-EM structure of the fly ribosome bound to dFMRP suggested that KH1 and KH2 domains interact near the peptidyl site (P site) of the 80S ribosome (*49*). Structural evidence that CHX binds to the tRNA exit site (E site) to block ribosome translocation may force the P site to remain fully occupied and unable to ratchet (*50, 51*). The growing nascent peptide bound to tRNA in the P site and CHX bound to the E site may occlude stable interaction between S499D FMRP and the ribosome.

The recovery levels observed in our FRAP assay was consistently around 20 ± 10% for FMRP reporters (Fig. 4). Although we have shown that the recovery was translation-dependent, a large population of FMRP did not recover, and their binding was translation-independent. The source of this unrecoverable population could be attributed to FMRP bound to the untranslated regions (UTRs) of mRNA. While there are differing opinions on whether FMRP can preferentially bind RNA in the coding sequence or the guanine-rich G-quadruplex structures on the 3′UTR (*10, 52*), our results may indicate the presence of both interactions. Moreover, if FMRP bound to CYFIP1 (*13*) or microRNA complexes (*14*) are present, a significant population of FMRP with longer binding-times may not recover in 20 minutes.

As FMRP is a negative translation regulator, our results support a clear partition of function for phosphorylation and dephosphorylation. Although FMRP-induced translation stalling has been simplified as an all-or-none event, it is likely that FMRP functions somewhere in between working in conjunction with other factors to regulate protein synthesis, such as mRNA stability through codon optimality (*47*), nonsense-mediated decay (NMD) (*53*) or m6A-mediated decay (*54*), all of which may not directly lead to or require completely stalled ribosomes. Moreover, it will be interesting to directly compare the translation rates of FMRP and non-FMRP target mRNAs to determine whether FMRP preferentially stalls or slows translation. In sum, our report demonstrates that FMRP granules are dynamic structures, and elucidating how FMRP asserts control of transport and local protein synthesis may benefit from high resolution studies.

## Methods

### Animals

Experimental procedures adhered to NIH and Albert Einstein College of Medicine Institutional Animal Care and Use Committee guidelines. Mice were group-housed in a standard 12 hr light/12 hr dark cycle. Dissociated mouse hippocampal neuron culture and acute transverse slices were prepared from male and female mice (P1 and P21-45, respectively): C57BL/6J (Charles River). Plasmids used in this work are available on Addgene.

### Plasmids and lentivirus production

Lentiviral expression vectors for GFP fused to FMRP, S499A and S499D were prepared by restriction digest cloning using PCR amplification of EGFP-FMRP sequence from p-EGFP-C1-Flag-mFmr1(wt), p-EGFP-C1-Flag-mFmr1(A) or p-EGFP-C1-Flag-mFmr1(D), respectively. The EGFP-FMRP, S499A and S499D plasmids were a gift from Stephanie Ceman (Addgene plasmids #87929, #87913 and #87914). HaloTag versions of FMRP reporters were generated by replacing GFP with HaloTag. The I304N mutant FMRP was generated using Q5 Site-Directed Mutagenesis Kit (NEB). G3BP1-GFP plasmid was a gift from Jeff Chao (Addgene plasmid #119950). Lentiviral particles were generated by transfecting ENV (pMD2.VSVG), packaging (pMDLg/pRRE), REV (pRSV-Rev) along with the expression vector into HEK293T cells using calcium phosphate. Viral supernatant was concentrated using Lenti-X concentrator (Takara Bio) according to manufacturer’s instruction. Virus particles were resuspended in Neurobasal A (Invitrogen) and stored at -80 °C. High-titer lentivirus were produced in the Einstein Genetic Engineering and Gene Therapy core according to standard protocol. Titer was quantified using fluorescence, RT-PCR and ELISA.

### Dissociated mouse hippocampal neuron culture

Post-natal day 1 (P1) mouse hippocampal tissue was isolated from C57BL/6 wildtype newborn pups. Hippocampi were dissociated in 0.25% trypsin, triturated and plated onto poly-D-lysine (Sigma) coated glass-bottomed dishes (MatTek) at 35,000-50,000 cells per dish. Neurons were cultured in Neurobasal A supplemented with B-27 (Invitrogen) and GlutaMAX (Invitrogen).

### Slice preparation

Acute transverse slices were prepared as follows: briefly, mice were decapitated, and brains were removed quickly and put into ice cold sucrose cutting solution or NMDG cutting solution. The sucrose cutting solution contained (in mM): 215 sucrose, 20 glucose, 26 NaHCO_3_, 4 MgCl_2_, 4 MgSO_4_, 1.6 NaH_2_PO_4_, 2.5 KCl, and 1 CaCl_2_. The NMDG cutting solution contained (in mM): 93 N-Methyl-d-glucamin, 2.5 KCl, 1.25 NaH_2_PO_4_, 30 NaHCO_3_, 20 HEPES, 25 D-glucose, 2 Thiourea, 5 Na-Ascorbate, 3 Na-Pyruvate, 0.5 CaCl_2_, 10 MgCl_2_. Mice over P35 were cut in NMDG. The hippocampi were isolated and cut using a VT1200S vibratome (Leica) at a thickness of 300 µm. The slices were then transferred to 32 °C ACSF for 30 minutes and then kept at room temperature (RT) for at least 1 hr before recording. For P21 mice: After ice-cold cutting, the slices recovered at RT (in 50% sucrose, 50% ACSF) for < 30 minutes and then at RT for 1 hr in ACSF. The artificial cerebrospinal fluid (ACSF) recording solution contained (in mM): 124 NaCl, 26 NaHCO_3_, 10 glucose, 2.5 KCl, 1 NaH_2_PO_4_, 2.5 CaCl_2_, and 1.3 MgSO_4_. All solutions were bubbled with 95% O_2_ and 5% CO_2_ for at least 30 minutes. All experiments in acute slices were performed at 25.5 ± 0.1 °C.

### Sample preparation and live imaging

Cultured hippocampal neurons were infected with lentivirus at DIV7 and imaged after 3 days at DIV10 or after one week at DIV14, unless noted otherwise. Prior to imaging, the neurobasal media was exchanged to Hiberate A low fluorescence (BrainBits) and allowed to equilibrate for 30 minutes at 37 °C prior to imaging at 22 °C. Cycloheximide (CHX, Sigma) 100 mg/ml or anisomycin (50 mM, Sigma) were introduced into the media after equilibration and allowed to incubate an additional 30 minutes at 37 °C prior to imaging at 22 °C. For HaloTag labeling, cells were treated with 10 nM JF549HTL or JF646HTL (Promega) for 15 minutes, followed by 3x washes and incubation in Hibernate A (low fluorescence) for at least one hour. Cells were washed again in Hibernate A prior to imaging at 35 °C. For single molecule imaging, 10 nM JFX549 and 10 nM JFX646 were mixed at 20:1 ratio and added to neuron cultures and labeled for 15 minutes and washed as above. The deuterated JFX Janelia Fluors were kindly provided by Dr. Luke Lavis at the Janelia Research Campus.

### Stereotaxic injections

WT C57BL/6J mice were stereotactically injected with high titer Halo-FMRP or Halo-S499A lentivirus into the dentate gyrus region between P21-P24 using established coordinates (-2.2 posterior to Bregma, 2.0 laterally, and 2.0 ventral from dura) using a total volume of 1.5 µL/hemisphere at a flow rate of 0.2 µL/min. After 2-3 weeks of recovery, mice were humanely killed and acute hippocampal slices were prepared as described above. For labeling Halo-FMRP, cell-permeable Halo-ligand was bath-applied. Slices were labeled with JF549-HTL (100 nM) in ACSF in a chamber oxygenated with 95%O_2_/5% CO_2_ for 1 hour. Slices were fixed with 4% PFA and mounted using ProLong Diamond (Invitrogen).

### Immunofluorescence

For immunofluorescence, neurons were fixed in 4% paraformaldehyde solution (PFA) and permeabilized. After blocking in normal goat serum, cells were treated with primary antibodies: anti-GFP (Aves lab) or anti-FMRP (Abcam; ab17722), followed by secondary antibodies: Goat anti-chicken Alexa Fluor 647 (ThermoFisher) or goat anti-rabbit Cy3 (Sigma) and DAPI. For hippocampal tissue, acute slices were washed twice in 1x PBS then incubated in blocking buffer (4% goat serum in 1x PBS + 2% BSA + 0.1% Tx-100) for 1 hr at RT. Primary antibodies were diluted directly into the antibody buffer (blocking buffer without goat serum) and floating slices were incubated overnight at 4 °C. After 4 washes with 1x PBS, slices were incubated in secondary antibodies (Invitrogen) diluted in blocking buffer overnight at 4 °C. Slices were washed 5x with 1x PBS, then mounted using ProLong Diamond (Invitrogen).

### Puromycylation

Acute slices were cut (as described in Slice preparation) 3 weeks after stereotaxic injection and checked for expression using an epifluorescence microscope. Slices were incubated with puromycin (50 µM, Sigma) for 30 minutes. Slices were then fixed for 1 hr in 4% PFA, blocked for 30 minutes, and stained according to the immunohistochemistry protocol above (primary antibodies: anti-ZnT3, rabbit polyclonal, 1:500, Synaptic Systems; anti-puromycin 1:1000, EMD Millipore) and imaged on the Zeiss LSM 880 with Airyscan using a LD LCI Plan-Apochromat 25x/0.8 mm Korr DIC M27 and 1.8x zoom. Images were Airyscan processed prior to analysis. Quantification was performed using FIJI by drawing regions of interest in the ZnT3 channel in which puromycin fluorescence intensity was measured in stratum lucidum. For stratum radiatum control, puromycin signal in regions of interest distal to the stratum lucidum as indicated by the ZnT3 labeling were measured.

### Fluorescence microscopy

Acquisition of fluorescence images were performed on a wide field microscope previously described (*55*). In brief, the inverted widefield microscope was equipped with 491 nm (Cobolt), 561 nm (Lasertechnik) and 640 nm (Coherent) laser lines along with a UApo 150x 1.45 NA or PlanApo 60x 1.45 NA oil immersion objectives (Olympus). Images were acquired on an EMCCD camera (Andor). Images were 512×512 in size with a pixel size of 106.7 nm (150x) or 266.7 nm (60x). Streaming videos of FMRP granules in dendrites were acquired by continuous imaging at 50 ms intervals for 400 or more frames from a single z-plane. All z-series images (0.2 mm steps) were maximum projected prior to analysis. For photobleaching fluorescent granules, we used the 405 nm UV laser (Omikron) along with a motorized focus lens (Thorlabs) which delivers focused light to a diffraction limited spot. After acquiring a pre-image, fluorescent granules were exposed to UV light at 10% power for 5 seconds for photobleaching. Typically, our photobleaching resulted in greater than 85% average photobleaching efficiency. Time-lapse z-series stacks (21 steps at 0.2 mm per step) were acquired for both 491 nm (GFP) and 561 nm (mCherry) channels every 30 seconds for 41 time-points (20 minutes). The cells were maintained at 22 °C throughout imaging through a humidified stage top incubator (Tokai Hit). For single molecule imaging of sparsely labeled Halo-FMRP or Halo-I304N, we first identified neurons labeled with JFX549 and switched to image JFX646. We acquired streaming images at 50 ms intervals for 1000 images (50 seconds) using 640 nm laser at high power (50%). We used the first 100 images for particle detection and tracking analyses. For tissue imaging, images were acquired on a Zeiss LSM 880 with Airyscan using a Plan-Apochromat 63x/1.4 NA oil DIC M27 and 1.8x zoom. Images were Airyscan processed prior to analysis. Threshold, laser power, and gain were kept constant for each experiment. Z-stacks of identical size were taken at similar depths were maximum projected.

### Image analysis

Analyses of imaging data were performed using FIJI (ImageJ). For particle detection and tracking we used the ImageJ plugin TrackMate (*56*). We used the Laplacian of Gaussian filter for detection where the [estimated blob diameter] was set to 5 pixels and [threshold] at 30. These values were determined empirically by evaluating several randomly selected reference images of GFP-FMRP and comparing the contrast values of all particles detected: [(intensity inside – intensity outside) / (intensity inside + intensity outside)](*56*). We tested a range of blob diameters from 3 to 6 pixel at one pixel intervals. Range of threshold values were tested from 15 to 50 at intervals of 5. From these conditions, we selected blob diameter and threshold values that yielded fewest particles with negative contrast values (where outside is brighter than inside of the puncta) while limiting detection of large spurious structures. We also used [median filter] and [sub-pixel localization] for improved gaussian fit and x-y localization of fluorescent puncta. These conditions were applied identically to all images to maintain consistency in the analysis. For analysis of Halo-FMRP and Halo-S499A, we used similar conditions for puncta analysis, except contrast since tissue generally resulted in variable background from slice to slice. To normalize, we used SNR (signal to noise) calculation in TrackMate: [(intensity inside – intensity outside) / standard deviation of the puncta intensity] (*56*). Statistical comparisons were made using unpaired, two-tailed student’s t-test or Mann-Whitney test in Prism.

For tracking granules, we used the default settings for the [Simple LAP] tracker with [maximum linking distance] of 5 pixels and [maximum gap-closing distance] of 5 pixels. In most situations puncta density did not result in overlapping particles. As such, we used the same value as our blob diameter as the upper-limit for non-diffusive motion. The [maximum gap-closing frame gap] was set to 2. To select for processively moving particles, we first identified all particle trajectories using TrackMate and sorted for processivity using our MATLAB script (available on github.com). In brief, each trajectory of a particle was rotated based on its longest axis of movement, which is the direction of processive movement. Once rotated, 5 or more consecutive positive or negative displacements in the long axis was defined as the processive movement. Kymographs of trafficking puncta were generated using the ImageJ plugin Kymographbuilder.

For single particle tracking, the [estimated blob diameter] was set to 5 pixels and [threshold] at 30. Under high illumination, single JFX dye molecules produce a point spread function that fits a gaussian profile within the 5-pixel diameter. Next, we used the [Simple LAP] tracker with [maximum linking distance] of 15 pixels and [maximum gap-closing distance] of 15 pixels, as the molecules were sparsely populated in the dendrites. The [maximum gap-closing frame gap] was set to 2. We used less stringent criteria to accommodate particles that were more confined in Halo-FMRP, relative to Halo-I304N particles that were more freely moving. In addition, we selected tracks with 5 or more continuously tracked frames to remove spurious detection of fast-moving dyes and shot noise from the detector.

For FRAP analysis, we selected a 9-pixel circular region of interest (ROI) around the bleach puncta and measured average intensity within the ROI for all timepoints. The average intensity values were normalized to pre image (t = -30 s) and the error bars denote standard deviation (SD). Background fluorescence was measured from a 14×14 area at least 100 pixels away from the site of photobleaching that did not contain any cellular fluorescence from all timepoints. The average background fluorescence was subtracted from the average fluorescence from the bleached ROI at each timepoint. To compare between drug treated and untreated FRAP experiments with different photobleaching efficiency, the first timepoint after photobleaching was set to baseline (zero) and all subsequent recovery values were baseline subtracted. The fluorescence recovery values were fit to a nonlinear regression curve (one-phase association) using Prism to derive time constants (τ) or time at half-maximal fluorescence of the recovery. The normalized fluorescence recovery at the final timepoint were analyzed using unpaired, two-tailed student’s t-test.

### Statistical analyses

In all results where statistical information was included, at least three or more independent trials were performed. Statistical comparisons were made using unpaired student’s t-test where p-values and sample sizes (n) were included in the Figure Legend. Standard deviation (SD) values were reported for photobleaching results and all others show standard error of the mean (SEM).

## Acknowledgments

We would like to thank Dr. Hannah Monday for critical reading and discussion of the manuscript. We also thank the Singer lab and Castillo lab members for their comments and discussions. We are grateful to the Lavis lab at HHMI/Janelia Research Campus for the JF dyes. We are also grateful to Stephanie Ceman and Jeff Chao for providing plasmids on Addgene. We would also like to acknowledge the shared instrument grant (S10OD025295) to Kostantin Dobrenis.

## Author contributions

Conceptualization: SCK, PEC, RHS, YJY

Methodology: SCK, PEC, YJY

Investigation: SCK, DH, YJY

Analysis: SCK, HC, KJY, YJY

Visualization: SCK, YJY

Supervision: PEC, RHS, YJY

Writing—original draft: SCK, YJY

Writing—review & editing: SCK, PEC, YJY

## Data and materials availability

All data needed to evaluate the conclusions in the paper are present in the paper and/or Supplementary Materials. Plasmids are available on Addgene. Code is available on GitHub.

