## Supplemental PDF for "Phosphorylation alters FMRP granules and determines their transport or protein synthesis abilities"

Shivani C. Kharod *et al.*

**This PDF file includes:**

Figs. S1 to S6
Movies S1 to S14

**Other Supplementary Materials for this manuscript include the following:**

Movies S1 to S14

**Supplementary figure 1**

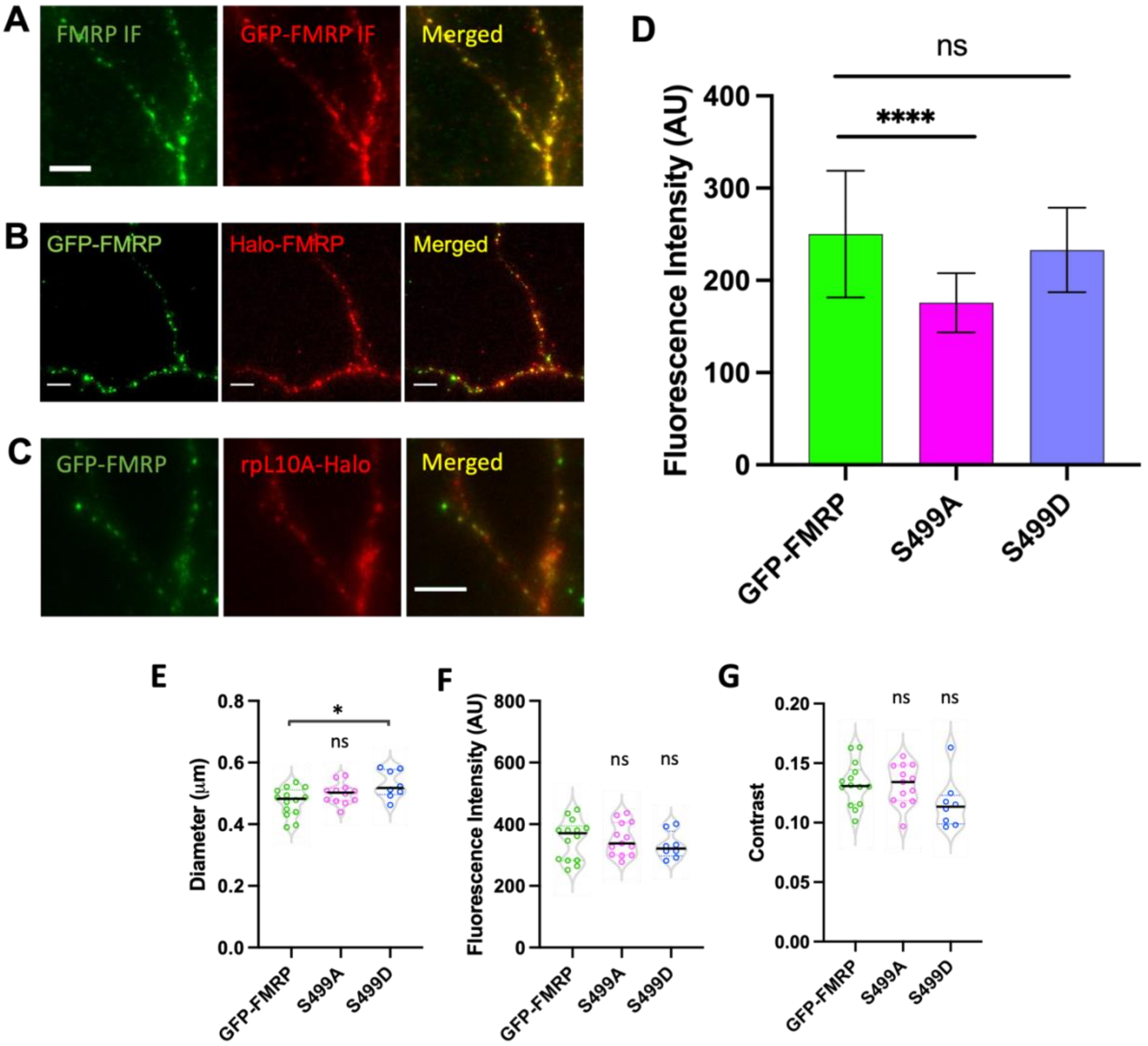


**Fig. S1.**

**FMRP reporters colocalize with endogenous FMRP and Halotagged ribosomal protein L10A.** (**A**) Immunofluorescence using antibodies to FMRP (Abcam; ab17722) shown in green and GFP (Aves lab) in red and merged image on the right. Cultured hippocampal neurons were infected with lentivirus encoding GFP-FMRP at DIV7, antibody labeled and imaged at DIV14. High degree of colocalization can be observed. Scale bar is 5 μm. (**B**) Live image of dendrites expressing both GFP-FMRP (green) and Halo-FMRP (red) labeled with JF646, along with the merged image on the right. Both reporters were expressed in neurons at DIV7 and imaged at DIV14. Many granules exhibit discrete colocalization of both reporters. Scale bar is 5 μm. (**C**) Live image of dendrites expressing GFP-FMRP (green) and ribosomal protein L10A-Halo (red) labeled with JF646, along with the merged image on the right. Many granules exhibit colocalization of reporter proteins. Both reporters were expressed in neurons at DIV7 and imaged at DIV14. Scale bar is 5 μm. (**D**) Average fluorescence intensities from dendrites expression GFP-FMRP (green), S499A (magenta) and S499D (blue). Statistical significance was calculated using unpaired, two-tailed student's t-test. $p^{****} < 0.0001$. (**E-G**) Violin plots of granule diameter, fluorescence intensity, or contrast from GFP-FMRP (green; n=14; 1466 granules), S499A (magenta; n=13; 1409 granules) and S499D (blue; n=8; 233 granules) after 3-days of expression. Horizontal lines denote population median. Statistical significance was calculated using unpaired, two-tailed student's t-test. $p^{*} < 0.05$; ns, not significant.

### Supplementary figure 2

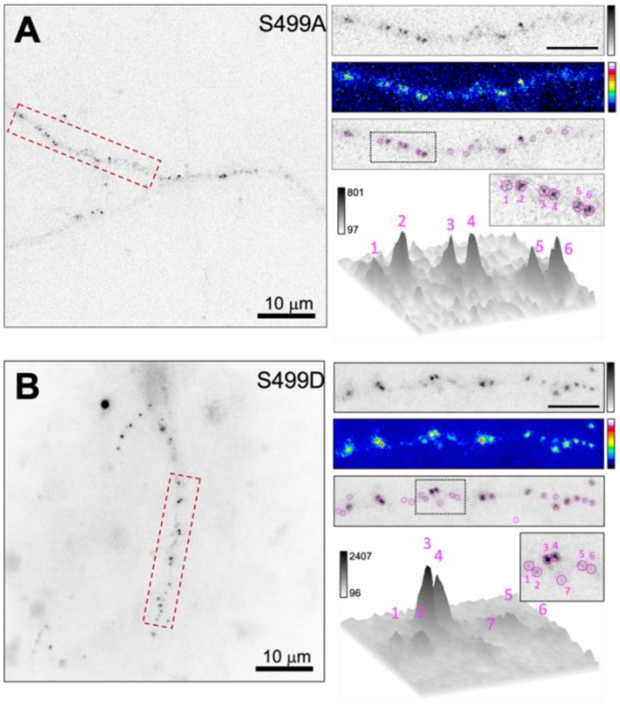


**Fig. S2.**

**S499 mutant FMRP reporters differ in their binding to FMRP granules.** (**A**) Representative dendrite expressing the GFP-S499A reporter (inverted grayscale). A dendritic segment is outlined in the dashed box. Scale bar is 10 µm. Top right panel shows the enlarged dendrite in the dashed box. Scale bar is 5 µm. Middle right panel shows the same dendrite in 16-color LUT. Bottom panel shows the dendrite overlaid with TrackMate particle detection (pink circles). Below the panel is a surface plot of S499A puncta shown in the dotted box. Numbers on surface plot correspond to puncta in inset. The LUT indicates the minimum and maximum fluorescence intensity range. (**B**) Representative dendrite expressing the GFP-S499D reporter (inverted grayscale). A dendritic segment is outlined in the dashed box. Scale bar is 10 µm. Top right panel shows the enlarged dendrite in the dashed box. Scale bar is 5 µm. Middle right panel shows the same dendrite in 16-color LUT. Bottom panel shows the dendrite overlaid with TrackMate particle detection (pink circles). Below the panel is a surface plot of S499D puncta shown in the dotted box. Numbers on surface plot correspond to puncta in inset. The LUT indicates the minimum and maximum fluorescence intensity range.

### Supplementary figure 3

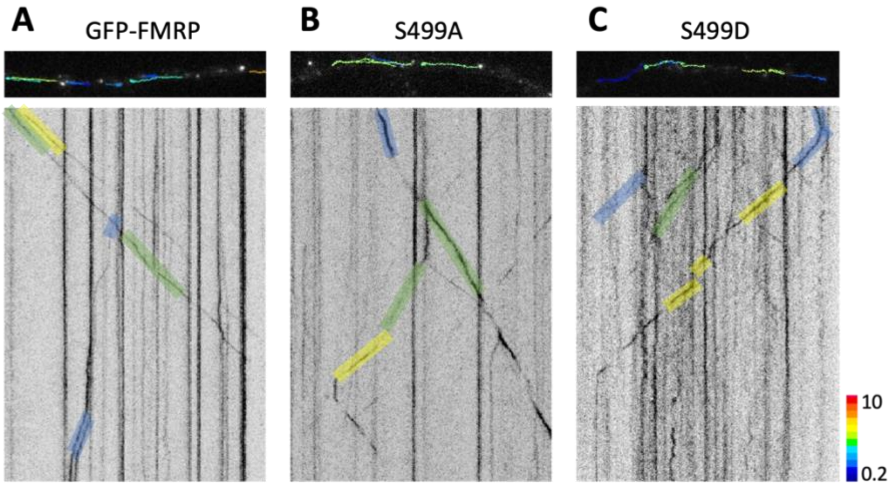


**Fig. S3.**

**FMRP reporter containing granules move along dendrites.** (**A**) Representative fluorescence image of GFP-FMRP dendrites overlaid with tracks detected by TrackMate. Dendrites were imaged at 50 ms per frame for 400 consecutive frames (20 seconds total; see also Supplementary movies S2 to S4). The streaming movies were converted to LUT-inverted kymograph to visualize moving particles along the x-t axis. Horizontal lines indicate non-moving particles and diagonal lines indicate moving particles where the slope indicate the speed (i.e. steeper slope is slower and flatter slope is faster). The tracks were overlaid with color-coded shades to indicate average speed of granule. (**B**) Fluorescence image of GFP-S499A expressing dendrite and kymograph. (**C**) Fluorescence image of GFP-S499D expressing dendrite and kymograph. LUT indicates average velocity of granules ranging from 0.2 to 10 μm/sec.

### Supplementary figure 4

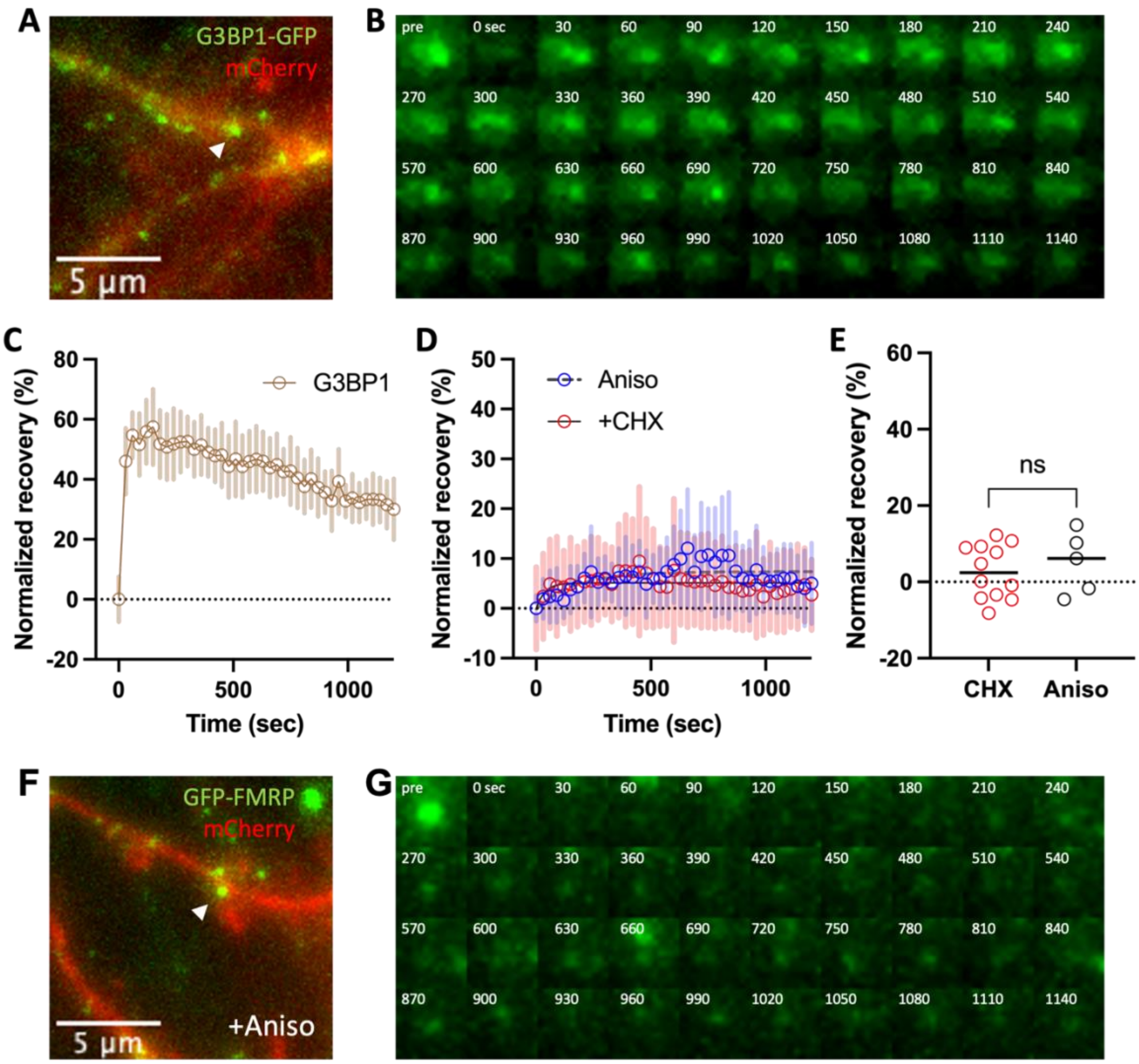


**Fig. S4.**

**FMRP granule recovery after photobleaching is different compared to G3BP1 and is sensitive to anisomycin.** (**A**) Representative image of G3BP1-GFP granule (green) in dendritic segment (red) selected for FRAP assay. White arrowhead indicates the photobleached granule and the scale bar is 5 μm. See also Supplementary movie S6. (**B**) Time-series montage of G3BP1-GFP granule recovery. Time is noted on the top-right in seconds. (**C**) Plot of normalized mean fluorescence recovery ± SD (shaded bars) for G3BP1-GFP granules (brown circles; n=9) over time. The dotted line indicates the average baseline intensity following photobleaching set to zero. (**D**) Plot of normalized average fluorescence recovery ± SD (shaded bars) for granules expressing GFP-FMRP under cycloheximide-treatment (CHX, red circles; n=12) or anisomycin-treatment (Aniso, blue circles; n=5) over time. The recovery values were fit to a nonlinear regression curve (gray dashed lines) to calculate time constants ($\tau$). The dotted line indicates the average baseline intensity following photobleaching set to zero. (**E**) Plot of normalized recovery at the final the timepoint for CHX-treatment (red circles; n=12) and Aniso-treatment (blue circles; n=5). The dotted line indicates the average baseline intensity following photobleaching set to zero. Horizontal lines denote population median. Statistical significance was calculated using unpaired, two-tailed student's t-test. ns, not significant. (**F**) Representative image of GFP-FMRP granule (green) in dendritic segment (red) selected for FRAP assay in the presence of anisomycin (+Aniso). White arrowhead indicates the photobleached granule and the scale bar is 5 μm. See also Supplementary movie S8. (**G**) Time-series montage of GFP-FMRP granule recovery in the presence of anisomycin. Time is noted on the top-right in seconds.

### Supplementary figure 5

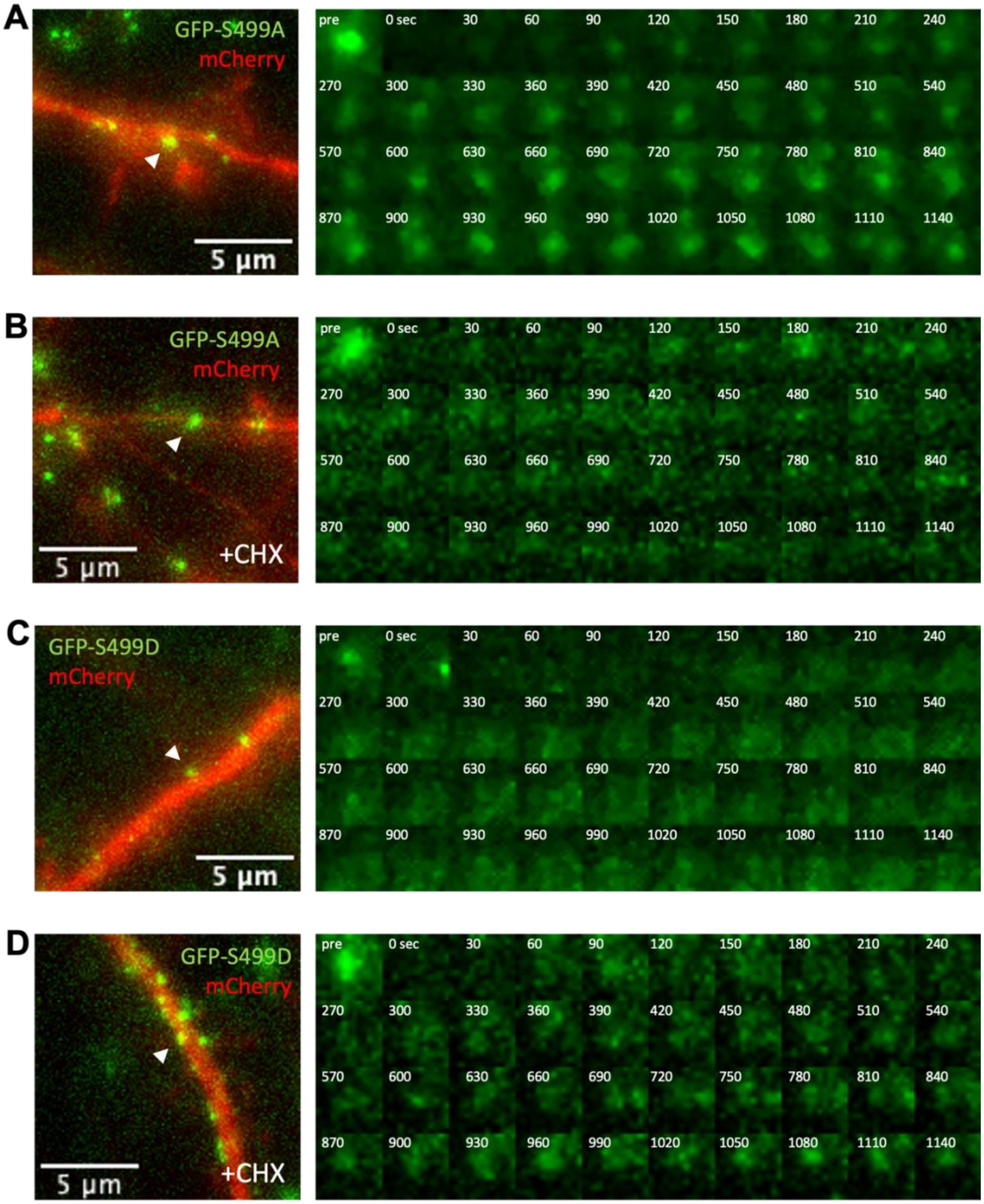


**Fig. S5.**
**FMRP phospho-mutant granule recovery after photobleaching differ between S499A and S499D mutants.** (**A**) Representative image of GFP-S499A granule (green) in dendritic segment (red) selected for FRAP assay. White arrowhead indicates the photobleached granule and the scale bar is 5 μm. See also Supplementary movie S9. On the right is a time-series montage of GFP-S499A granule recovery. Time is noted on the top-right in seconds. (**B**) Representative image of GFP-S499A granule (green) in dendritic segment (red) selected for FRAP assay in the presence of

cycloheximide (+CHX). White arrowhead indicates the photobleached granule and the scale bar is 5 mm. See also Supplementary movie S10. On the right is a time-series montage of GFP-S499A granule recovery. Time is noted on the top-right in seconds. (**C**) Representative image of GFP-S499D granule (green) in dendritic segment (red) selected for FRAP assay. White arrowhead indicates the photobleached granule and the scale bar is 5 μm. See also Supplementary movie S11. On the right is a time-series montage of GFP-S499D granule recovery. Time is noted on the top-right in seconds. (**D**) Representative image of GFP-S499D granule (green) in dendritic segment (red) selected for FRAP assay in the presence of cycloheximide (+CHX). White arrowhead indicates the photobleached granule and the scale bar is 5 μm. See also Supplementary movie S12. On the right is a time-series montage of GFP-S499D granule recovery. Time is noted on the top-right in seconds.

**Supplementary figure 6**

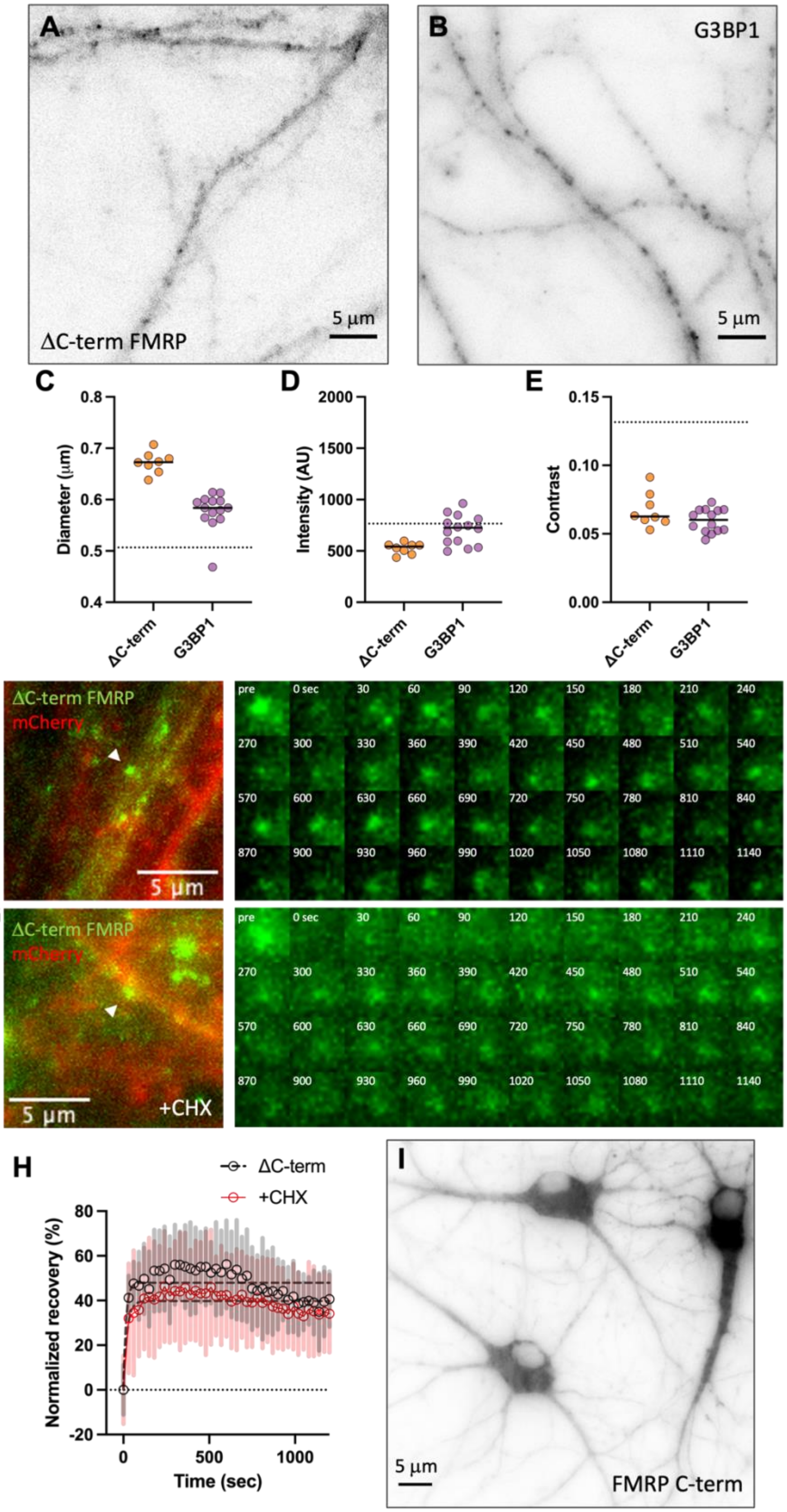

**Fig. S6.**
**S499 is necessary but not sufficient for granule binding.** (**A**) Representative fluorescence image (pseudo-colored as inverted grayscale) of a dendrite expressing the mutant FMRP reporter that does not have the C-terminal region (ΔC-term). Scale bar is 5 μm. (**B**) Representative image of G3BP1-GFP granules in dendrites. Scale bar is 5 μm. (**C**) Scatter plot of average diameter of fluorescent granules in neurons expressing ΔC-term FMRP (orange; n=8; 2768 granules) or G3BP1 (purple; n=14; 3422 granules). Horizontal lines denote population median. Dashed line indicates median GFP-FMRP granule diameter (0.507 ± 0.008 μm). (**D**) Scatter plot of average intensity of fluorescent granules in neurons expressing ΔC-term FMRP (orange; n=8) or G3BP1 (purple; n=14). Horizontal lines denote population median. Dashed line indicates median GFP-FMRP granule intensity (765.3 ± 54.5 AU). (**E**) Scatter plot of average contrast of fluorescent granules in neurons expressing ΔC-term FMRP (orange; n=8) or G3BP1 (purple; n=14). Horizontal lines denote population median. Dashed line indicates median GFP-FMRP granule contrast (0.132 ± 0.005). (**F**) Representative image of ΔC-term FMRP granule (green) in dendritic segment (red) selected for FRAP assay. White arrowhead indicates the photobleached granule and the scale bar is 5 μm. See also Supplementary movie S13. On the right is a time-series montage of ΔC-term FMRP granule recovery. Time is noted on the top-right in seconds. (**G**) Representative image of ΔC-term FMRP granule (green) in dendritic segment (red) selected for FRAP assay in the presence of cycloheximide (+CHX). White arrowhead indicates the photobleached granule and the scale bar is 5 μm. See also Supplementary movie S14. On the right is a time-series montage of ΔC-term FMRP granule recovery. Time is noted on the top-right in seconds. (**H**) Plot of normalized average fluorescence recovery ± SD (shaded bars) for ΔC-term FMRP granules (black circles; n=8) and cycloheximide-treatment (CHX, red circles; n=8) over time. The recovery values were fit to a nonlinear regression curve (gray dashed lines) to calculate time constants (τ). The dotted line indicates the average baseline intensity following photobleaching set to zero. (**I**) Representative fluorescence image (pseudo-colored as inverted grayscale) of cultured neurons expressing the FMRP C-terminal region (FMRP C-term). Scale bar is 5 μm.

**Movie S1.**
**Single particle tracking of Halo-FMRP and Halo-I304N.** On the top-left is the Halo-FMRP raw images and on the top-right is the Halo-FMRP particles detected by TrackMate (pink circles) overlaid with trajectories color-coded by total displacement. On the bottom-left is the Halo-I304N raw images and on the bottom-right is the Halo-FMRP particles detected by TrackMate (pink circles) overlaid with trajectories color-coded by total displacement. The movie is running at 50 frames per second.

**Movie S2.**
**GFP-FMRP granules in dendrites overlaid with tracks.** On the left is a dendrite expressing GFP-FMRP and on the right is the same dendrite overlaid with particle tracks detected by TrackMate color-coded by displacement. The movie is running at 50 frames per second.

**Movie S3.**
**GFP-S499A granules in dendrites overlaid with tracks.** On the left is a dendrite expressing GFP-S499A and on the right is the same dendrite overlaid with particle tracks detected by TrackMate color-coded by displacement. The movie is running at 50 frames per second.

**Movie S4.**
**GFP-S499D granules in dendrites overlaid with tracks.** On the left is a dendrite expressing GFP-S499D and on the right is the same dendrite overlaid with particle tracks detected by TrackMate color-coded by displacement. The movie is running at 50 frames per second.

**Movie S5.**
**GFP-FMRP granule recovery after photobleaching.** On the left is a dendrite expressing GFP-FMRP and mCherry where the photobleached granule is circled. Shown on the right is GFP-FMRP only. The movie is running at 15 frames per second.

**Movie S6.**
**G3BP1-GFP granule recovery after photobleaching.** On the left is a dendrite expressing G3BP1-GFP and mCherry where the photobleached granule is circled. Shown on the right is G3BP1-GFP only. The movie is running at 15 frames per second.

**Movie S7.**
**GFP-FMRP granule recovery after photobleaching in cycloheximide.** On the left is a dendrite expressing GFP-FMRP and mCherry where the photobleached granule is circled. Shown on the right is GFP-FMRP only. The movie is running at 15 frames per second.

**Movie S8.**
**GFP-FMRP granule recovery after photobleaching in anisomycin.** On the left is a dendrite expressing GFP-FMRP and mCherry where the photobleached granule is circled. Shown on the right is GFP-FMRP only. The movie is running at 15 frames per second.

**Movie S9.**
**GFP-S499A granule recovery after photobleaching.** On the left is a dendrite expressing GFP-S499A and mCherry where the photobleached granule is circled. Shown on the right is GFP-S499A only. The movie is running at 15 frames per second.

**Movie S10.**
**GFP-S499A granule recovery after photobleaching in cycloheximide.** On the left is a dendrite expressing GFP-S499A and mCherry where the photobleached granule is circled. Shown on the right is GFP-S499A only. The movie is running at 15 frames per second.

**Movie S11.**
**GFP-S499D granule recovery after photobleaching.** On the left is a dendrite expressing GFP-S499D and mCherry where the photobleached granule is circled. Shown on the right is GFP-S499D only. The movie is running at 15 frames per second.

**Movie S12.**
**GFP-S499D granule recovery after photobleaching in cycloheximide.** On the left is a dendrite expressing GFP-S499D and mCherry where the photobleached granule is circled. Shown on the right is GFP-S499D only. The movie is running at 15 frames per second.

**Movie S13.**
**ΔC-term FMRP granule recovery after photobleaching.** On the left is a dendrite expressing ΔC-term FMRP and mCherry where the photobleached granule is circled. Shown on the right is ΔC-term FMRP only. The movie is running at 15 frames per second.

**Movie S14.**
**ΔC-term FMRP granule recovery after photobleaching in cycloheximide.** On the left is a dendrite expressing ΔC-term FMRP and mCherry where the photobleached granule is circled. Shown on the right is ΔC-term FMRP only. The movie is running at 15 frames per second.
